# Cas-CLIP: a method for customizing pooled CRISPR libraries

**DOI:** 10.1101/187757

**Authors:** Jiyeon Kweon, Da-eun Kim, An-Hee Jang, Yongsub Kim

**Author notes:** These authors contributed equally to this work (Jiyeon Kweon, Da-eun Kim). Correspondence should be addressed to Y. K.

## Abstract

Although pooled CRISPR libraries are widely used in high-throughput screening to study various biological processes, library construction for researcher’s own study is a time-consuming, labor-intensive, and expensive process. In this study, we develop a simple, scalable method, called Cas-CLIP, to customize conventional pooled CRISPR libraries using the CRISPR/Cas9 system. We show that conventional pooled CRISPR libraries can be modified by eliminating gRNAs that target positive genes, enabling the identification of unknown target genes in CRISPR screening. Cas-CLIP is a precise method for customizing conventional pooled CRISPR libraries and will broaden the scope of high-throughput screening technology.

## INTRODUCTION

Various pooled libraries, such as shRNA and cDNA libraries, have been developed for high-throughput functional studies(Boehm and Hahn, 2011). Currently, the Clustered regularly interspaced short palindromic repeat (CRISPR)-Cas9 technology for genome engineering is evolving rapidly(Kim and Kim, 2014) and CRISPR-based pooled libraries are widely used in high-throughput functional studies to understand various biological processes(Shalem et al., 2015). The pooled CRISPR library, comprising guide RNAs (gRNAs) targeting individual genes in the genome, is used with Cas9 or Cas9 variants in loss-of-function or gain-of-function studies(Gilbert et al., 2014; Hess et al., 2016; Klann et al., 2017; Konermann et al., 2015; Shalem et al., 2014; Wang et al., 2014). Although several groups provide pooled CRISPR libraries via the non-profit company, Addgene, researchers still require customized libraries for their investigations(Kurata et al., 2016). Since the generation of new libraries for researcher’s own study is a time-consuming, labor-intensive, and expensive process, we developed an innovative method, called Cas-CLIP (CRISPR/Cas-based library customization from pooled CRISPR library), for customizing conventional pooled CRISPR libraries. We show that the sequence-specific depletion of desired gRNAs from conventional CRISPR libraries, without potential off-target effects, is possible and these ‘CLIPed’ libraries can be used for high-throughput screening. Furthermore, we demonstrate that 81 gRNAs, targeting 27 kinase genes related to the targets of FDA-approved drugs can be simultaneously eliminated from pooled CRISPR libraries. Based on our results, Cas-CLIP can be employed to customize conventional pooled CRISPR libraries for use in various biological screening procedures.

## RESULTS

### Development of the Cas-CLIP method to customize conventional CRISPR libraries

We first noted that gRNAs in the pooled CRISPR library are expressed from the U6 promoter (originating from human or mouse) and the promoter contains the sequence 5′-CCG-3′ upstream of the gRNA-encoding region. Using these as PAM sequences for Cas9 proteins, reverse-complementary gRNAs (rc-gRNAs) could be designed to target each gRNA in the pooled CRISPR plasmid DNA library, which would eliminate specific gRNAs from the pooled CRISPR library (Figure 1). As proof of concept, we performed Cas-CLIP to remove *HPRT1* gRNAs from the human GeCKOv2 library(Sanjana et al., 2014). Cas9 ribonucleoproteins (RNPs), comprising Cas9 proteins and *in vitro*- transcribed rc-gRNAs targeting *HPRT1* gRNAs, were treated to the GeCKOv2 plasmids DNA library *in vitro*.

**Figure 1.**
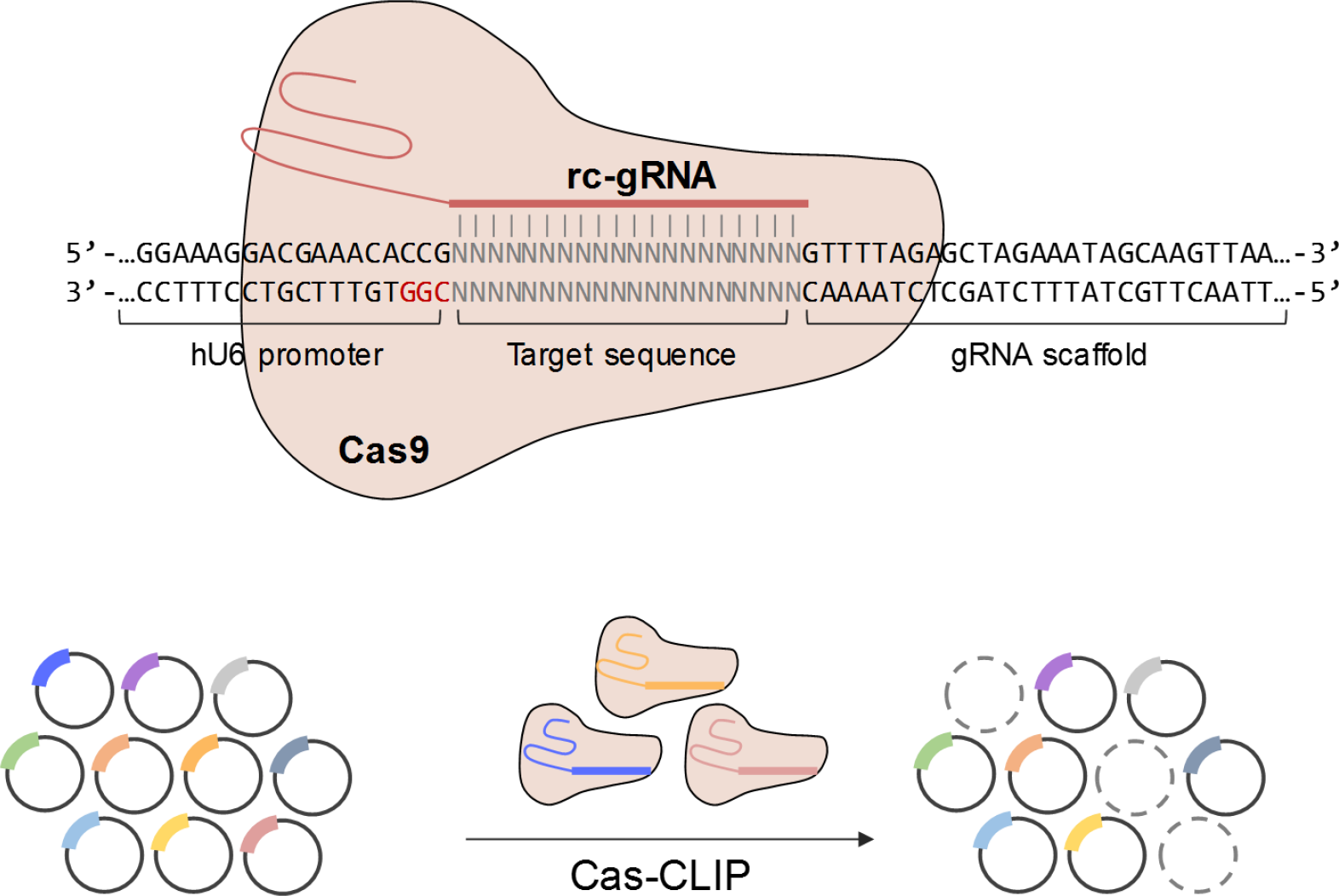
Schematic outline of Cas-CLIP. The gRNA-encoding plasmid DNA in the pooled CRISPR library contained the nucleotide sequence 5’-CGG-3’, which could be used as the PAM sequence for Cas9 proteins. Cas9 RNPs, which are complexes of Cas9 proteins and rc-gRNAs, could selectively cleave plasmid DNA possessing complementary sequences with rc-gRNA. Based on this principle, only the desired gRNAs could be depleted from the pooled CRISPR library.

To confirm *HPRT1* gRNA cleavage, we performed polymerase chain reaction (PCR) or quantitative PCR (qPCR) using target-specific primers (Figure 2a). While using *HPRT1*- target site-specific primer pairs, PCR was conducted successfully with wild-type GeCKOv2 libraries; however, we could not detect PCR amplicons with ‘*HPRT1*-CLIPed’ GeCKOv2 libraries (Figure 2b). We confirmed that the gRNAs were depleted dose- and time-dependently (Figure S1). Additionally, we performed targeted deep sequencing to verify *HPRT1* gRNA depletion. Among 123,411 gRNAs in total (65,383 in GeCKOv2 A library and 58,028 in GeCKOv2 B library), only six *HPRT1* gRNAs were dramatically depleted in the ‘*HPRT1*-CLIPed’ GeCKOv2 libraries (Figures 2c, d and Table S1). Moreover, Cas-CLIP was performed with another pooled CRISPR library type, GeCKOv1 and by targeted deep sequencing; 1 to 7 gRNAs were confirmed to be successfully depleted from the wild type library (Figure 2 and Table S2), showing that Cas-CLIP could selectively remove only plasmids of interest from pooled CRISPR libraries.

**Figure 2.**
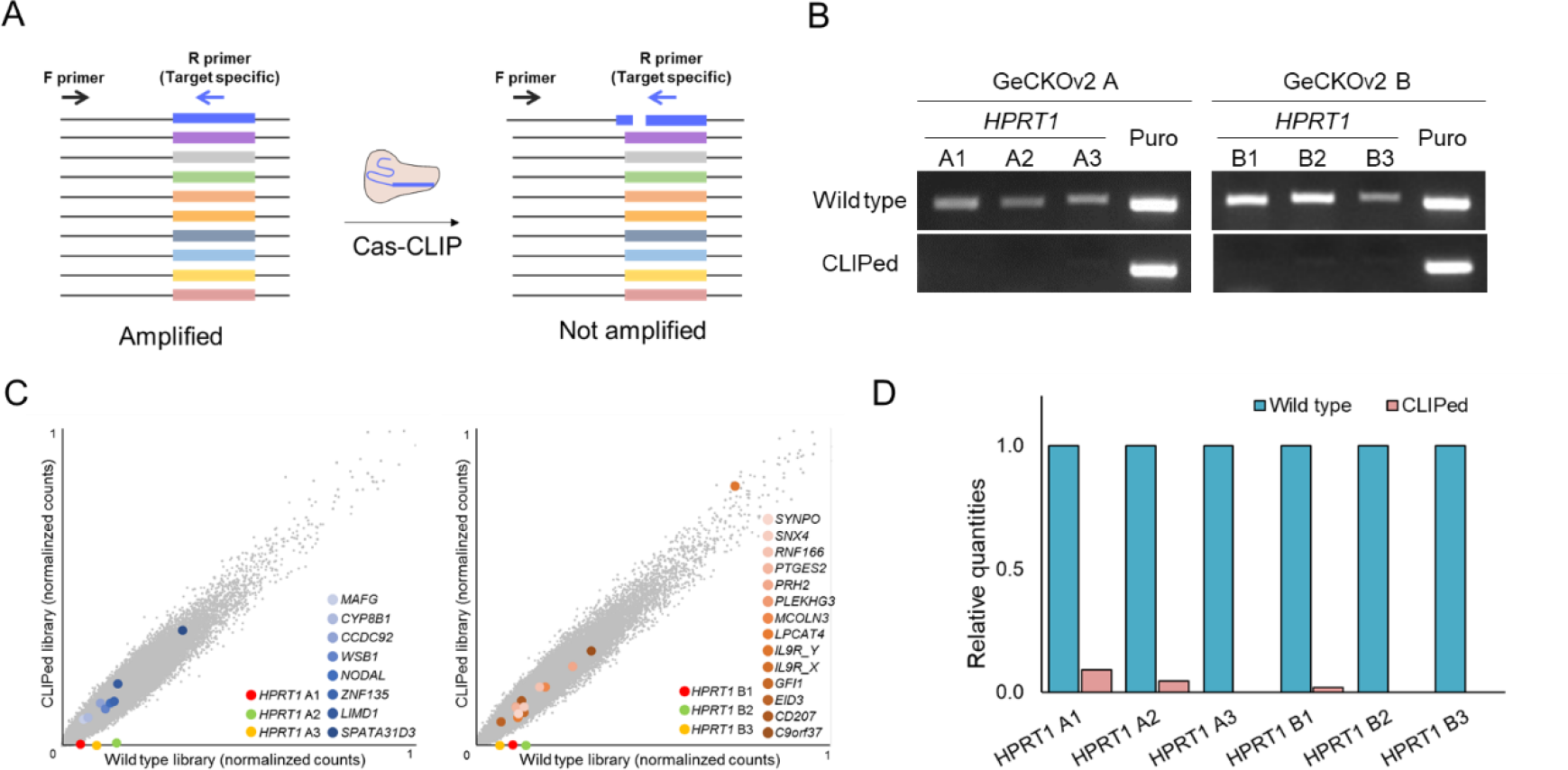
Cas-CLIP for ‘*HPRT1*-CLIPed’ CRISPR library generation. **(a)** Schematic outline of PCR to assess the quality of ‘CLIPed’ library. The forward primer annealed to the U6 promoter and the reverse primers could selectively anneal to the target sequences. While using the primer pairs, amplification occurred in the wild-type library, although not in the ‘CLIPed’ library. **(b)** The PCR amplicons of the ‘*HPRT1*-CLIPed’ GeCKOv2 library were analyzed by agarose gel electrophoresis. Each of the GeCKOv2 A and B libraries had three *HPRT1* gRNAs, all of which were depleted from each library. For all six reverse primers annealed to the *HPRT1* gRNA-encoding plasmid DNA, amplification occurred with the wild-type libraries as templates, although not with the ‘*HPRT1*-CLIPed’ libraries. A pair of primers annealing to puromycin-encoding sequences in the plasmid DNA was used for internal control amplification. The primer sequences are listed in Table S5. **(c)** Targeted deep sequencing analysis of ‘*HPRT1*-CLIPed’ libraries as described previously(Kweon et al., 2017). Scatter plots of ‘*HPRT1*-CLIPed’ GeCKOv2 A (left) and ‘*HPRT1*-CLIPed’ GeCKOv2 B (right) libraries showed that only *HPRT1* gRNAs were depleted from the wild-type GeCKOv2 libraries. **(d)** Relative *HPRT1* gRNA quantities were calculated from the targeted deep sequencing data. Numerical data are presented in Table S1.

### Potential off-target effects of Cas-CLIP

We carefully considered the potential off-target effects of Cas-CLIP. Since all plasmid DNA in the pooled CRISPR libraries had the same PAM sequence, we assumed that the potential off-target effect could be determined only by the mismatch tolerance of rc-gRNAs. Generally, pooled CRISPR libraries are designed to minimize potential off-target effects on the genome, in order that the gRNA nucleotide sequences contained therein are not similar(Shalem et al., 2014). Additionally, more than 3-bp mismatches between gRNA and target sequence can be distinguished by Cas9 nucleases, which can inhibit the off-target effects(Cho et al., 2014). We examined whether there were gRNAs with sequences similar to those of the six *HPRT1* gRNAs in the GeCKOv2 libraries and in fact, there was 1 gRNA with 5-bp mismatches and 21 gRNAs with 6-bp mismatches with the gRNAs (Table S3). The relative quantities of the above-mentioned gRNAs in the wild-type and ‘*HPRT1*-CLIPed’ libraries did not differ significantly from each other (Figure 2c and Table S3). These results showed that Cas-CLIP could be successfully applied to conventional CRISPR libraries and the potential off-target effect of Cas-CLIP was not of concern while using well-defined pooled CRISPR libraries.

### High-throughput screening with CLIPed libraries

To determine whether the ‘CLIPed’ library could be used for functional studies, we performed 6-thioguanine (6-TG) screening with the wild-type and ‘*HPRT1*-CLIPed’ libraries. As 6-TG is cytotoxic to *HPRT1*-expressing cells, *HPRT1* gRNAs result in positive hits in 6-TG screening because they disrupt *HPRT1* and make cells resistant to 6-TG(Kim et al., 2017; Koike-Yusa et al., 2014). Lentiviral particles were produced from the wild-type and ‘*HPRT1*-CLIPed’ libraries and transduced into a HeLa-Cas9 stable cell line. To measure relative quantities of gRNAs in the infected cells, genomic DNA was isolated from these cells and analyzed by targeted deep sequencing at a depth of over 80-fold. Similar to the findings on plasmid DNA, of 123,411 gRNAs, only *HPRT1* gRNAs were dramatically depleted in two independent experiments (Figure 3a and Table S1). After confirming that the gRNAs in the cells was sufficient coverage, we conducted recessive screening with 6-TG and observed that 6-TG-resistant clones were not present in the cells infected with the ‘*HPRT1*-CLIPed’ libraries (Figure 3b). The targeted deep sequencing analysis revealed that all *HPRT1* gRNAs were highly enriched in the cells infected with the wild-type library, although not in those infected with the ‘*HPRT1*-CLIPed’ library, as in a previous study(Kim et al., 2017) (Figures 3c and S3 and Table S1). Our results showed that Cas-CLIP could successfully eliminate gRNAs of interest *in vitro* and the ‘CLIPed’ library could be used for high-throughput functional studies.

**Figure 3.**
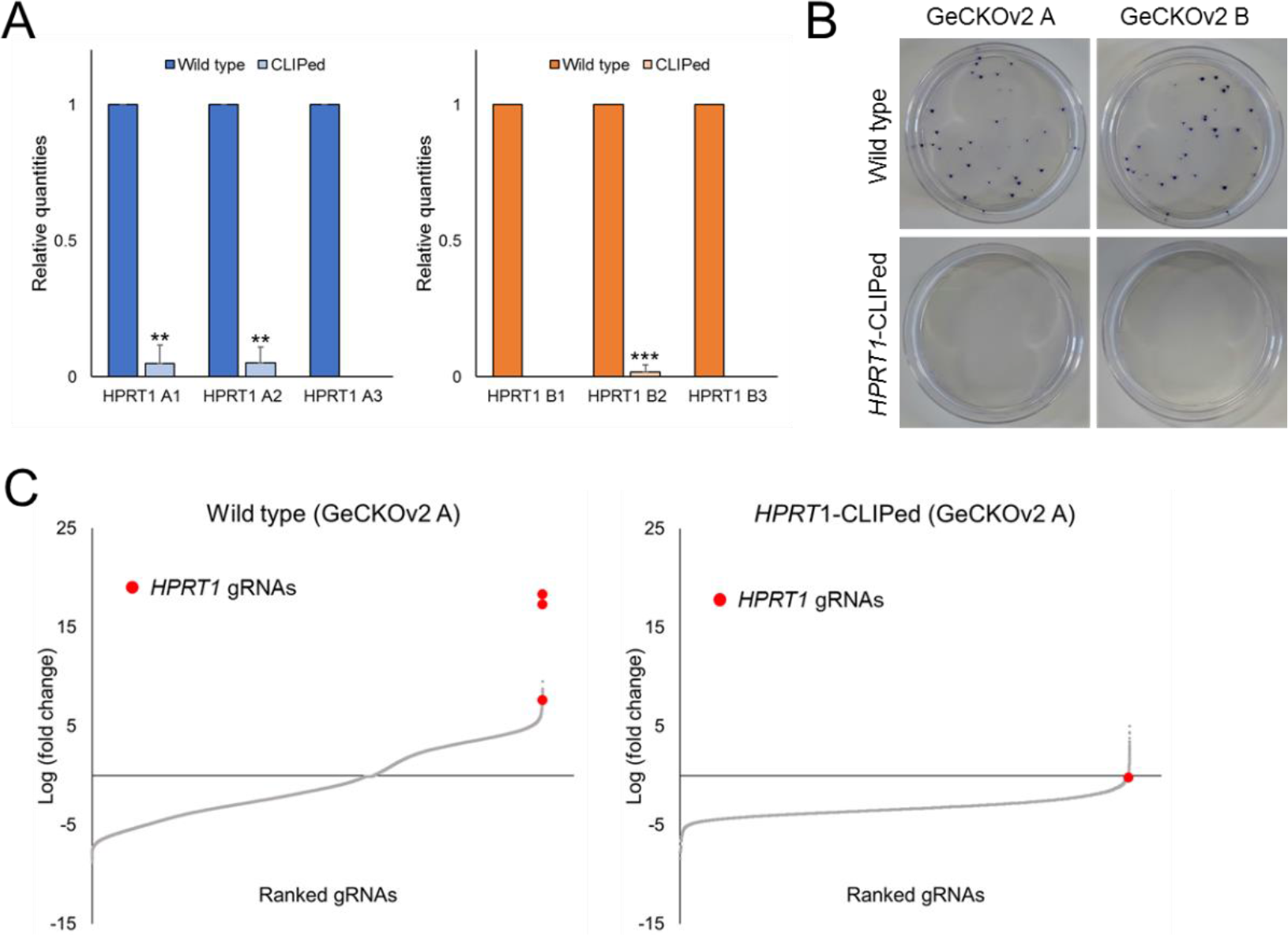
High-throughput screening with ‘*HPRT1*-CLIPed’ libraries. *(a)* Relative *HPRT1* gRNA quantities were calculated from the targeted deep sequencing data on the cells infected with each library. Error bars indicate SEM. \*\**P* < 0.01, \*\*\**P* < 0.001. Student’s *t*-test. **(b)** Crystal violet staining images after 6-TG selection of cells infected with designated libraries. 6-TG-resistant clones were observed only in the cells infected with the wild-type libraries. **(c)** Log_2_-fold change in gRNAs in cells infected with the wild-type or ‘HPRT1-CLIPed’ GeCKOv2 A library after 6-TG selection. All three *HPRT1* gRNAs were highly enriched in the cells infected with the wild-type library. Numerical data are presented in Table S1.

### Depletion of multiple gRNAs with CLIP

Scalability of Cas-CLIP is essential for applications in various biological research studies. To confirm that a set of gRNAs targeting each gene could be simultaneously depleted by Cas-CLIP, we designed rc-gRNAs for 81 gRNAs targeting 27 human kinase genes, which were known to be the targets of FDA-approved drugs(Xu et al., 2016). Kinases are known to be potential therapeutic targets and their inhibitors are used as cancer-targeting agents(Wu et al., 2015). However, recently, non-kinase targets of kinase inhibitors have been identified and they are gaining attention as new therapeutic targets(Munoz, 2017). To construct a ‘kinase-CLIPed’ library, 81 rc-gRNAs, which were simultaneously transcribed *in vitro*, and Cas9 proteins were treated together to GeCKOv2 library A as described below in the Methods section and the quality of the ‘kinase-CLIPed library’ was assessed by targeted deep sequencing. The relative quantities of gRNAs showed that all 81 gRNAs in the ‘kinase-CLIPed’ library were dramatically depleted compared with those in the wild-type library (Figures 4a, b and Table S4). This ‘kinase-CLIPed’ library may be used for biomedical screening to identify new druggable target genes. Taken together, we show that Cas-CLIP could simultaneously deplete multiple gRNAs from conventional CRISPR libraries without off-target effects and the ‘CLIPed’ libraries could be used for high-throughput screening.

**Figure 4.**
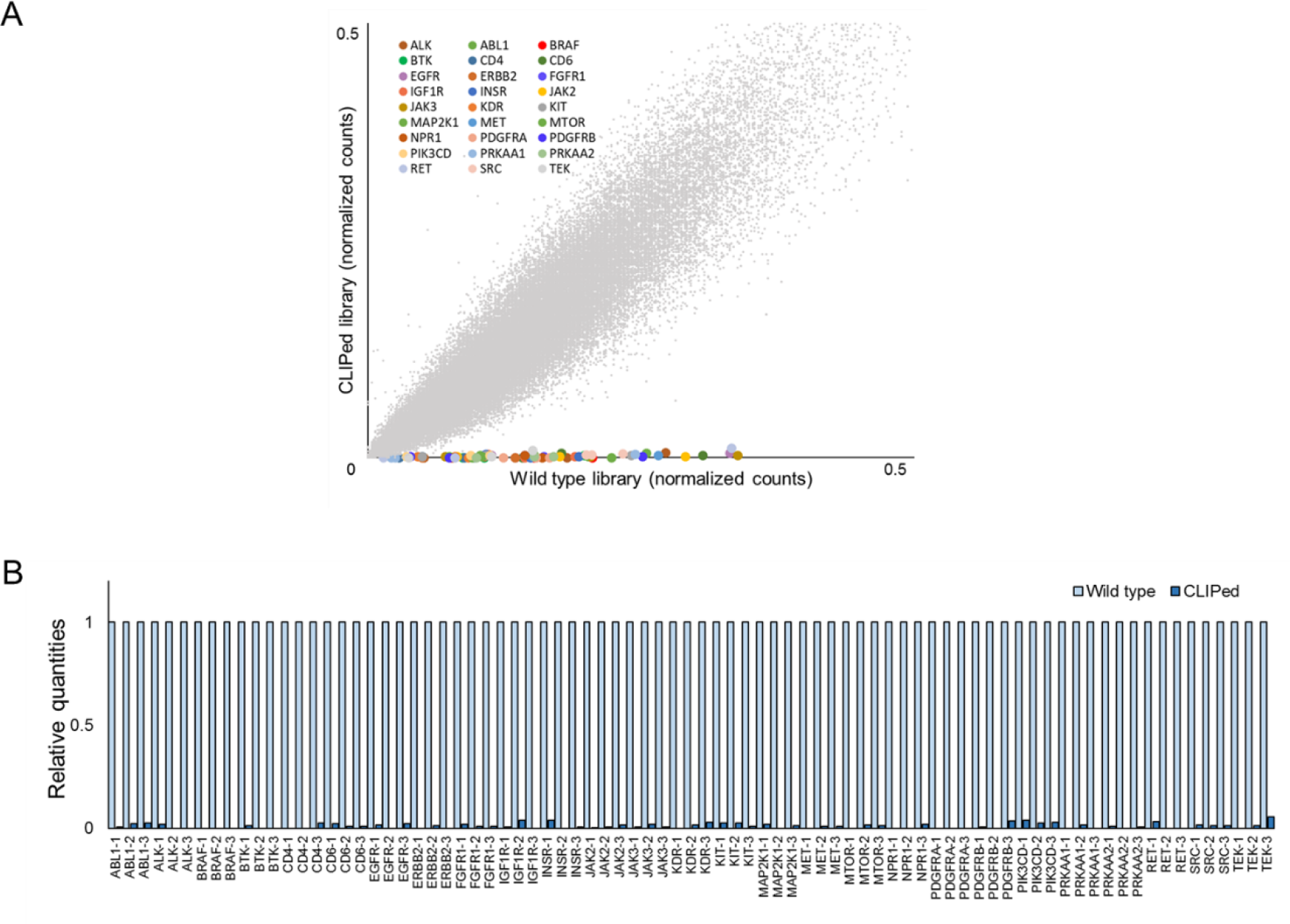
Cas-CLIP scalability for depletion of multiple gRNAs. **(a)** Targeted deep sequencing analysis of ‘kinase-CLIPed’ GeCKOv2 A library. The scatter plots showed that all 81 gRNAs that targeted 27 human kinase genes were depleted in the ‘kinase-CLIPed library’ compared with the wild-type library. **(b)** Relative gRNA quantities were calculated from the deep sequencing data. Numerical data are presented in Table S4.

## DISCUSSION

We developed a method named Cas-CLIP, which is a precise and scalable tool for customizing conventional pooled CRISPR libraries. Although previous studies show that Cas9 RNP can be used for eliminating unwanted DNAs such as mitochondrial rRNA in RNA-Seq libraries(Gu et al., 2016; Wu et al., 2016), we first showed that Cas-CLIP could deplete only gRNAs of interest and multiple gRNAs could simultaneously be eliminated in pooled CRISPR libraries. The advantage of Cas-CLIP over the generation of new libraries is its ability to reduce the time to construct a pooled library (Figure S4). Furthermore, we presume that Cas-CLIP can reduce cost and labour while removing up to 100 gRNAs (Figure S5). Interestingly, the deep sequencing results showed HPRT1 gRNAs remained in the ‘CLIPed’ library at low frequencies, but the HPRT1 gRNAs were not enriched after 6-TG screenings using the ‘CLIPed’ library. Based on these results, it appeared that the HPRT1 gRNAs, interpreted as remaining in the ‘CLIPed’ libraries, were either due to deep sequencing errors or did not exceed the threshold that could affect library screening.

Cas-CLIP can be applied to genetic screening in diverse ways, for example, to exclude positive genes or gene sets in recessive screening for reducing the number of genetic screening rounds required to identify unknown target genes. Other types of Cas proteins which have different types of PAM sequences, such as Cpf1 with the T-rich PAM sequences, can be adopted to extend the usefulness of Cas-CLIP. Furthermore, Cas- CLIP is applicable to other library types, such as shRNA and cDNA libraries, with the potential to expand the availability of various pooled libraries in biomedical research.

## METHODS

### Purification of Cas9 proteins and *in vitro* transcription of rc-gRNAs

Recombinant Cas9 proteins were expressed in *Escherichia coli* and purified by affinity chromatography using Ni-NTA agarose beads. LB broth (400 mL) containing 50 µg/mL kanamycin was inoculated with 4 mL of overnight-cultured BL21 (DE3) cells containing pET-His6-spCas9 plasmid DNA. When optical density at 600 nm (OD_600_) reached 0.6, Cas9 protein expression was induced by treating the cells with 0.2 mM IPTG for 16 h at 18°C. The cell pellet was harvested by centrifugation at 6,000 × g for 20 min and stored at −80°C until purification. For protein purification, the cell pellet was thawed on ice for 15 min and lysed with 4 mL of lysis buffer (50 mM NaH_2_PO_4_, 300 mM NaCl and 10 mM imidazole, pH 8.0) supplemented with 1 mM PMSF, 1 mM DTT, 1 mg/mL lysozyme and 600 units of Benzonase^®^ Nuclease. The cell lysate was cleared by centrifugation at 10,000 × g for 30 min and applied to Ni-NTA agarose beads according to the manufacturer’s instructions and an imidazole gradient (20 mM wash buffer and 250 mM elution buffer) were used to wash and elute the proteins. The eluent was buffer-exchanged with Cas9 storage buffer (20 mM HEPES, 150 mM KCl, 1 mM DTT and 40% glycerol, pH 7.5) using a PD-10 desalting column and concentrated using Amicon^®^ Ultra-4 100K according to the manufacturer’s instructions. The purified Cas9 proteins were separated by SDS-PAGE. gRNAs were synthesized *in vitro* using T7 RNA polymerase. Templates for gRNA synthesis were synthesised by annealing and extending two complementary oligonucleotides, which are listed in Table S5. For a 100-µL *in vitro* transcription reaction, T7 RNA polymerase buffer containing 1.5 µg of PCR products, each NTP at 4 mM, 14 mM MgCl_2_ and 100 units of RNase inhibitor was incubated with 500 units of T7 RNA polymerase for 3 h at 37°C. Subsequently, the reaction mixture was treated with DNase I for 30 min at 37°C and the *in* vitro-transcribed gRNAs were purified using an miRNeasy kit.

### Cas-CLIP for gRNA depletion from pooled CRISPR libraries

For generating the ‘CLIPed’ library, 20 µg of pooled CRISPR library plasmids DNA (human GeCKO v1 and v2 lentiviral sgRNA libraries, Addgene #1000000049) was incubated with NEBuffer 3.1 containing 1× Cas9 RNPs and a complex of 10 µg of Cas9 proteins and 7.5 µg of rc-gRNAs for 8 h at 37°C. Subsequently, the mixture was treated with RNase A for 30 min at 37°C and ‘CLIPed’ libraries were purified by isopropanol precipitation.

### qPCR and targeted deep sequencing

qPCR and targeted deep sequencing were performed to assess the quality of the ‘CLIPed’ libraries. For qPCR, a common forward primer, annealing to the U6 promoter, and target-specific reverse primers were used with the 2× iQ™ SYBR^®^ Green Supermix. A primer pair specific to the puromycin gene in the plasmid DNA was used for internal control PCR amplifications. One nanogram of pooled libraries was subjected to qPCR and the comparative C_T_ method was used to estimate specific gRNA depletion. For targeted deep sequencing, PCR amplicons of gRNA-encoding regions were sequenced on MiniSeq or NextSeq system as described previously(Kweon et al., 2017). Sequencing data were analyzed using count_space.py as described previously(Joung et al., 2017).

### Cell culture and CRISPR library screening

HEK293T cells (ATCC CRL-11268) and HeLa cells (ATCC, CCL-2) were maintained in Dulbecco’s modified Eagle’s medium (DMEM) supplemented with 10% fetal bovine serum (FBS) and 1% penicillin/streptomycin. HEK293T cells (5 × 10^6^) were seeded on a 100-mm dish one day before transfection and transfected with 15 µg of the lentiviral library and two viral packaging plasmids (9 µg of psPAX2 and 6 µg of pMD2.G) using Lipofectamine 2000 according to the manufacturer’s instructions and the culture medium was changed after 6 h. The lentiviral particles were harvested and filtered using a 0.45-µm filter 48 h after transfection.

HeLa cells (1 × 10^7^) were plated on a 100-mm dish and transduced with the lentivirus (MOI ∼0.1). The transduced cells were selected on media containing 1 µg/mL puromycin. For 6-TG screening, 1 µg/mL 6-TG was added to the culture medium 3 days after transduction. After two weeks, crystal violet staining was performed to visualize 6-TG-resistant clones and genomic DNA was isolated for analysis by targeted deep sequencing. The screening data was analysed using the pipeline MAGeCK (ver. 0.5.6)(Li et al., 2014)

## Data availability

The data supporting the observations of this study are available from the corresponding author upon reasonable request. All targeted deep sequencing data were deposited at the Sequence Reads Archive database of the NCBI with accession number SRP133450.

## ACKNOWLEDGEMENTS

We thank Heon Seok Kim for the helpful discussion of this study. This study was supported by the National Research Foundation of Korea (2016R1D1A1A02937096, 2017M3A9B4062419, and 2016R1A6A3A04009014).

## AUTHOR CONTRIBUTIONS

J.K., D.K. and Y.K. designed the study and J.K., D.K. and A.H.K. performed the experiments. Y.K. supervised the study. All authors discussed the results and commented on the manuscript.

